# Revisiting the STRmix^™^ likelihood ratio probability interval coverage considering multiple factors

**DOI:** 10.1101/2021.06.25.449960

**Authors:** Jo-Anne Bright, Shan-I Lee, John Buckleton, Duncan Taylor

**Affiliations:** ESR, Private Bag 92021, Auckland, New Zealand; Department of Statistics, University of Auckland, New Zealand; School of Biological Sciences, Flinders University, GPO Box 2100, Adelaide, SA 5001, Australia; Forensic Science SA, PO Box 2790, Adelaide, SA 5000, Australia

## Abstract

In previously reported work a method for applying a lower bound to the variation induced by the Monte Carlo effect was trialled. This is implemented in the widely used probabilistic genotyping system, STRmix^™^. The approach did not give the desired 99% coverage.

However, the method for assigning the lower bound to the MCMC variability is only one of a number of layers of conservativism applied in a typical application. We tested all but one of these sources of variability collectively and term the result the near global coverage. The near global coverage for all tested samples was greater than 99.5% for inclusionary average *LR*s of known donors. This suggests that when included in the probability interval method the other layers of conservativism are more than adequate to compensate for the intermittent underperformance of the MCMC variability component. Running for extended MCMC accepts was also shown to result in improved precision.

## Introduction

We have previously reported [1] the underperformance of the method (called the highest posterior density, HPD) used in STRmix^™^ to assign a lower bound to the variability caused by the MCMC process. This under performance was demonstrated by measuring the coverage of the lower bound, termed here the MCMC coverage. It applied to only the MCMC uncertainty component of the HPD in isolation, however this is not how HPD intervals are usually calculated in practice. In practice the HPD lower bound will typically also include allele frequency variability. This uncertainty is inherent to sampling a subset of individuals from a population in order to form an allele frequency database. It is also possible within a HPD interval calculation in STRmix^™^ to include uncertainty in the value assigned to the co-ancestry within a population (F_ST_, or theta, *θ*), done by supplying theta as a distribution. However, more typical in practice is to assign a fixed co-ancestry value that is in the higher plausible range of theta for the population being considered. These two additional aspects within the final reported HPD lower bound *LR* add conservatism to the value. It is also possible to add layers of conservatism to the reported *LR*, outside of the HPD, for example by reporting a unified *LR* [2, 3] or truncating the estimate.

There has been concern from the community regarding the results of [1] that the *LR* value reported is no longer representative of a 99% lower bound, or worse the results have been misunderstood to mean that the bound being reported is no longer conservative at all (personal communication). In truth, the addition of the MCMC uncertainty aspects within the HPD only increased the conservatism of the reported interval, and we have always maintained that when considered within the context of all other aspects of conservatism built into the reported value, it would still attain greater than 99% coverage. Because there are multiple layers of conservativism used in STRmix^™^ we attempt here a test of all but one of these layers collectively. We term this the near global coverage (NGC). In the NGC we seek to measure the variability within the MCMC component, uncertainty in allele probabilities, and the behaviour of the population genetic model.

The MCMC coverage has previously been reported [1].

To assess coverage inclusive of the allele frequencies, we take a set of frequencies and treat them as “true probabilities” only to create the “true *LR*.” Each lower bound estimate is created using a set of frequencies that has been resampled with replacement from these data, and hence will differ slightly from the truth and from each other.

The population genetic model used in STRmix^™^ is based on NRC II recommendation 4.2 [4, 5]. Previous work suggests that this population genetic model when used with the actual correct value for *θ* is conservative [6, 7]. At *θ*= 0 the model becomes the product rule. The product rule has been shown to have a mild bias in the non-conservative direction. In Figure 1 we give a graphical representation of these behaviours from data similar to that reported in Curran et al. [7] (hereafter CTB). CTB give the results of an experiment measuring the magnitude of the subpopulation effect.

**Figure 1.**
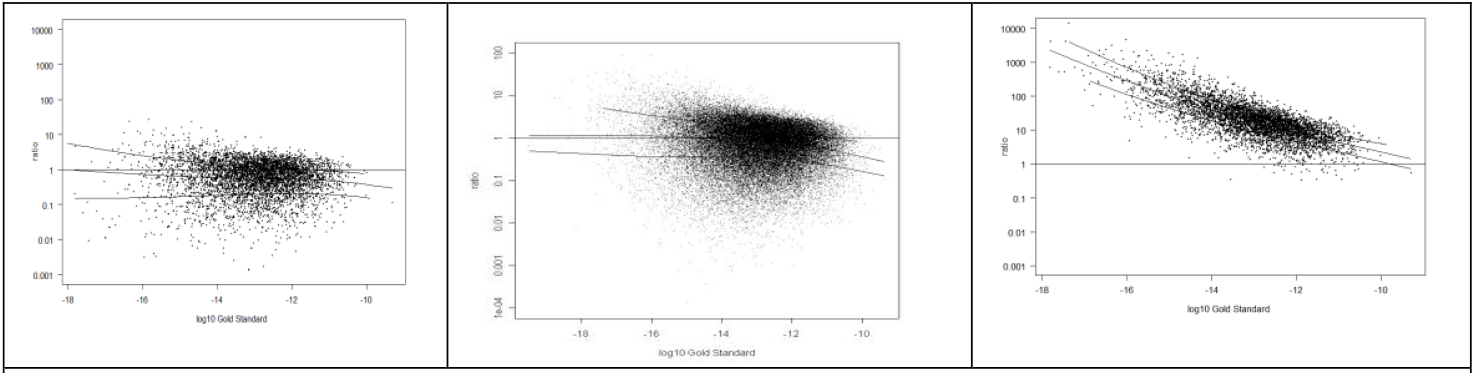
The ratio of the estimated genotype frequency to the true frequency for the product rule (left) NRC II recommendation 4.1 (centre) and NRC II recommendation 4.2 (right). These figures were developed from data constructed in the way described in CTB. The true *θ* created in the simulation is 0.03 and this is the value used in recommendations 4.1 and 4.2.

Curran et al. [7] give the measured performances that are reproduced in Table 1.

**Table 1.** The percentage of non-conservative estimates given by the product rule and NRC recommendation 4.2 for *θ* = 0.01 and *θ* = 0.03 from Curran et al. [7].

| Model | Number of subpopulations | Not conservative |  |
| --- | --- | --- | --- |
| | | $\theta = 0.01$ | $\theta = 0.03$ |
| Product rule | 2 | 48.2% | 65.1% |
|  | 10 | 51.5% | 64.8% |
|  | 50 | 50.9% | 59.9% |
| NRC II recommendation 4.2 | 2 | 1.6% | 0.8% |
|  | 10 | 0.5% | 0.6% |
|  | 50 | 0.1% | 0.1% |

An α % lower bound is intended to be constructed so that α % of the time the true value is higher than the bound. It has been suggested that if the process was rerun there could be other lower bounds and that some lower bounds could be even lower than the one quoted. This is self-evident. It is not possible for each lower bound to be lower than all the others. However, it is not the relationship of the lower bounds to each other that is sought but rather their relationship to the true answer.

## Method

In previously reported work 11 profiles were deconvoluted *N* = 1000 times each within STRmix^™^ V2.7 [8, 9]. All profiles were interpreted with eight chains each of 10,000 burn-in accepts and 50,000 post burn-in accepts. All profiles were GlobalFiler^™^ (Thermo Fisher CA) separated on a 3500 capillary electrophoresis instrument. Seven of the profiles were from the PROVEDIt dataset (two single-source, three two-person mixtures, and two three-person mixtures) and two of the profiles were simulated based on peak height variances observed across the PROVEDIt dataset (one two-person and one three-person mixture) [10]. The remaining two profiles (one single-source and one two-person mixture) were selected because one known contributor gave an *LR* supporting exclusion due to significant dropout across the profile. These deconvolutions also form the basis of this study.

To assess coverage inclusive of the allele frequencies we take a set of frequencies, in this case the FBI Caucasian frequencies [11], and treat them as the truth. We use these “true probabilities” only to create the “true *LR*”. Each lower bound estimate is created using a set of frequencies that has been resampled with replacement from these data, and hence will differ slightly from the truth and from each other.

To test the conservativeness induced by the use of a conservative value for theta in NRC II recommendation 4.2 we in this study use a fixed value of theta, applied outside the HPD method. We assume that the use of *θ* = 0.005 is close to the mean of the plausible range for theta for Caucasians. This seems reasonable from the data published in Buckleton et al. [12] and Steele and Balding [13]. We term this the near global ground truth (NGGT). The CTB data suggests that if the value for theta is approximately correct the model is still conservative about 99% of the time. We do not assess the effect of this last layer of conservativism as that would require a large simulation. We may attempt this in the future.

In this work, for the coverage test inclusive of MCMC, allele probability, and conservative value for theta (NGC) we create the MCMC, allele probability, and fixed theta value “true” *LR* (NGGT) by taking the log of the average of the sub-source *N* point estimates. This was estimated using the FBI Caucasian frequencies with *θ* = 0.005 (NGGT).

As discussed earlier, *θ* = 0.005 is a plausible value for informing an approximately correct value for theta, however the *LR* subsequently calculated using the population genetic model in use for STRmix^™^ is still likely to be conservative. The word “true” is placed in quotations because we recognize that we do not know the true value, that the value we are using is still likely to be conservative, and that we are assigning this based on plausible values from published studies.

The lower bounds were created inclusive of allele frequency and MCMC uncertainty, using a typically conservatively chosen theta, in this case *θ* = 0.01, and varying, externally from STRmix^™^, the allele probabilities using numerical resampling from the original database (FBI Caucasian allele frequencies [11]). Hence, we have introduced the effect of allele probability variability, MCMC variability, and a typically used value for *θ* into the lower bounds. As per the previous study, for each deconvolution this was repeated 1000 times and the lower 0.99 quantile taken as the lower α = 0.99 bound.

Each of the *N* lower bounds were scored as to whether it was below or above NGGT. This gives the NGC.

## Results

A typical result for two of the MCMC, allele probability, and subpopulation effect coverage tests featuring *LR*s around 10^2^ to 10^4.5^ are given in Figures 2 and 3. The points in Figure 2 and 3 represent the 99% lower bound HPD intervals when calculated inclusive of allele frequency and MCMC uncertainty, *θ* = 0.01 and resampled allele frequency databases. Each point not only represents a different sampled database, but also a different one of the 1000 deconvolutions. The distribution of these HPD intervals is shown to the right of the points, along with the point which corresponds to the upper 99% of the values (grey line). If exactly 99% coverage were achieved, then the NGGT (indicated by a blue dashed line in Figures 2 or 3) would exactly align with the grey line on the distribution. If they fall below the grey line, then they have not achieved 99% coverage and if they fall above the line then they have exceeded 99% coverage.

**Figure 2.**
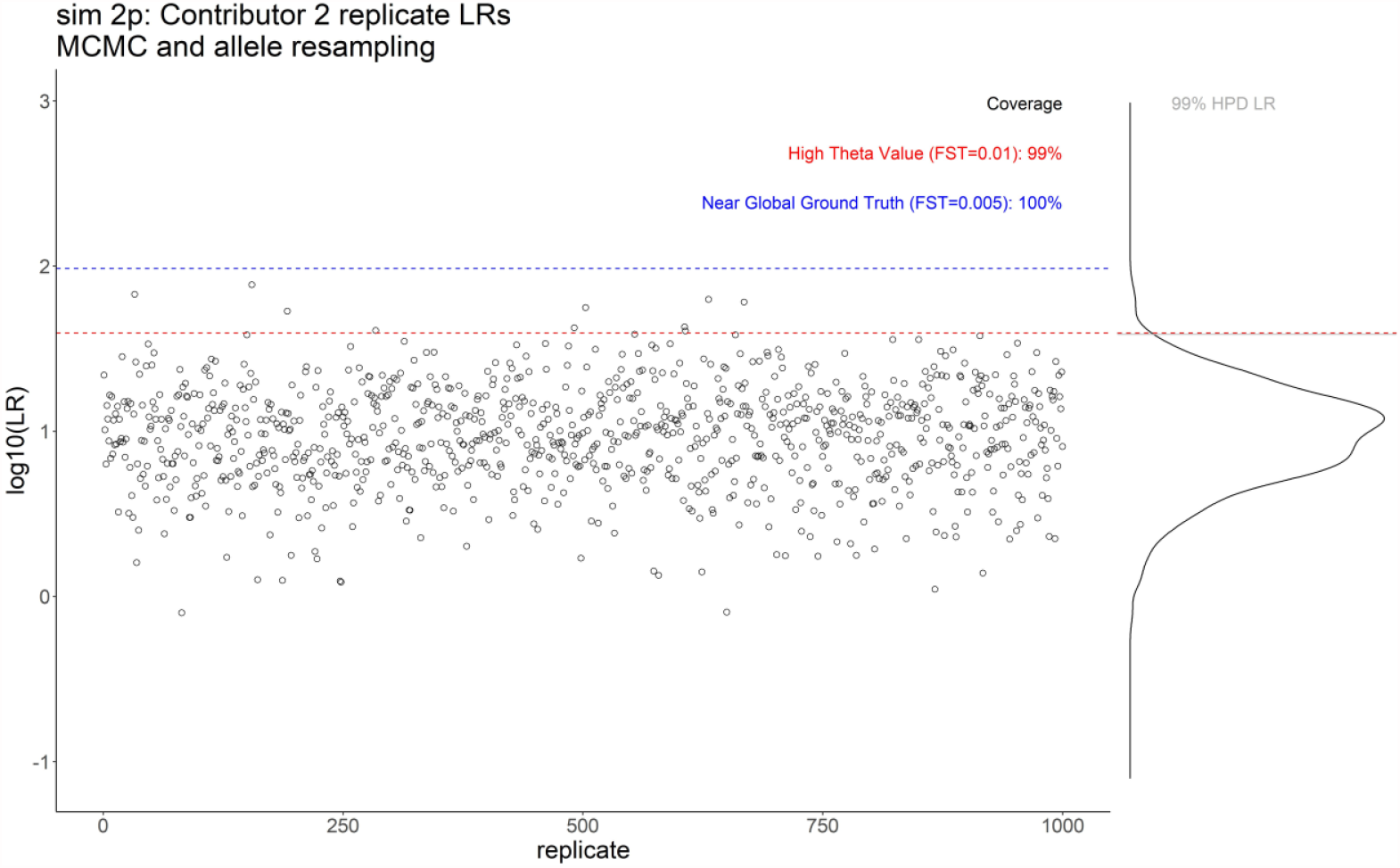
Comparison of each of 1000 replicates for the sample sim 2p contributor reference 2 comparison. The individual log_10_*LR* (α = 99% lower bound inclusive of both MCMC and allele probability resampling, with *θ* = 0.01) are given as hollow black circles. Also shown are HTV as a red dashed line and NGGT as a blue dashed line. At the right-hand margin these data are given as a probability density distribution, and the upper 99% quantile of the HPD density distribution is marked as a grey line on the distribution. The percentage coverage with respect to each ground truth *LR* is provided in the legend.

**Figure 3.**
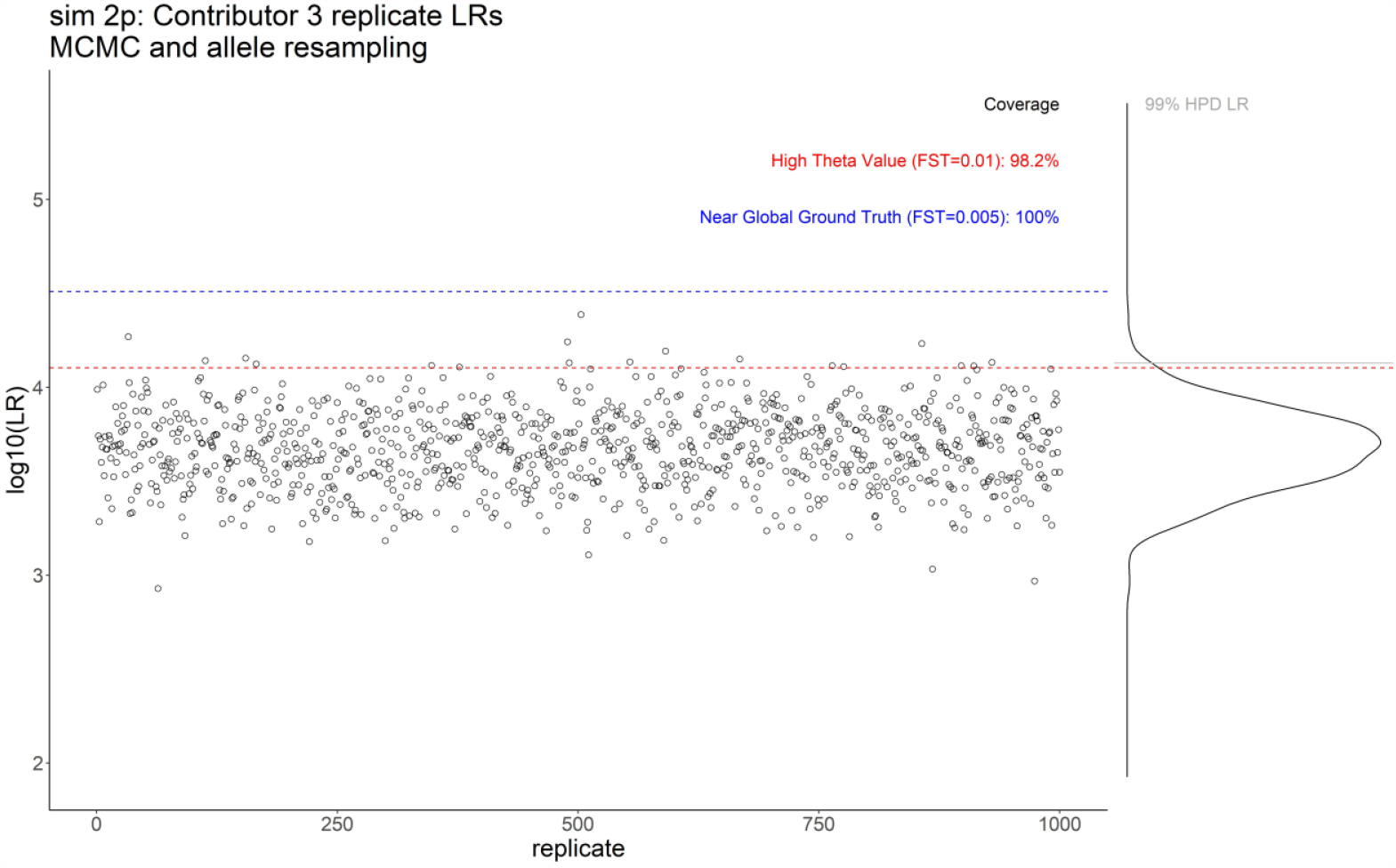
Comparison of each of 1000 replicates for the sample sim 2p contributor reference 3 comparison. The individual log_10_*LR* (α = 99% lower bound inclusive of both MCMC and allele probability resampling, with *θ* = 0.01) are given as hollow black circles. Also shown are HTV as a red dashed line and NGGT as a blue dashed line. At the right-hand margin these data are given as a probability density distribution, and the upper 99% quantile of the HPD density distribution is marked as a grey line on the distribution. The percentage coverage with respect to each ground truth *LR* is provided in the legend.

Also shown is the *LR* that would occur using the ground truth from the Monte Carlo and allele probabilities but using a high theta – red dashed line, termed high theta *LR* value (HTV).

The result for the three-person mixture, A08 3p contributor reference 3 with *LR*s around 10^32^, is shown in Figure 4. The previously reported coverage [1] considering only MCMC variation was the lowest observed (76.1%). The plots for all other comparisons are given in the Appendix.

**Figure 4.**
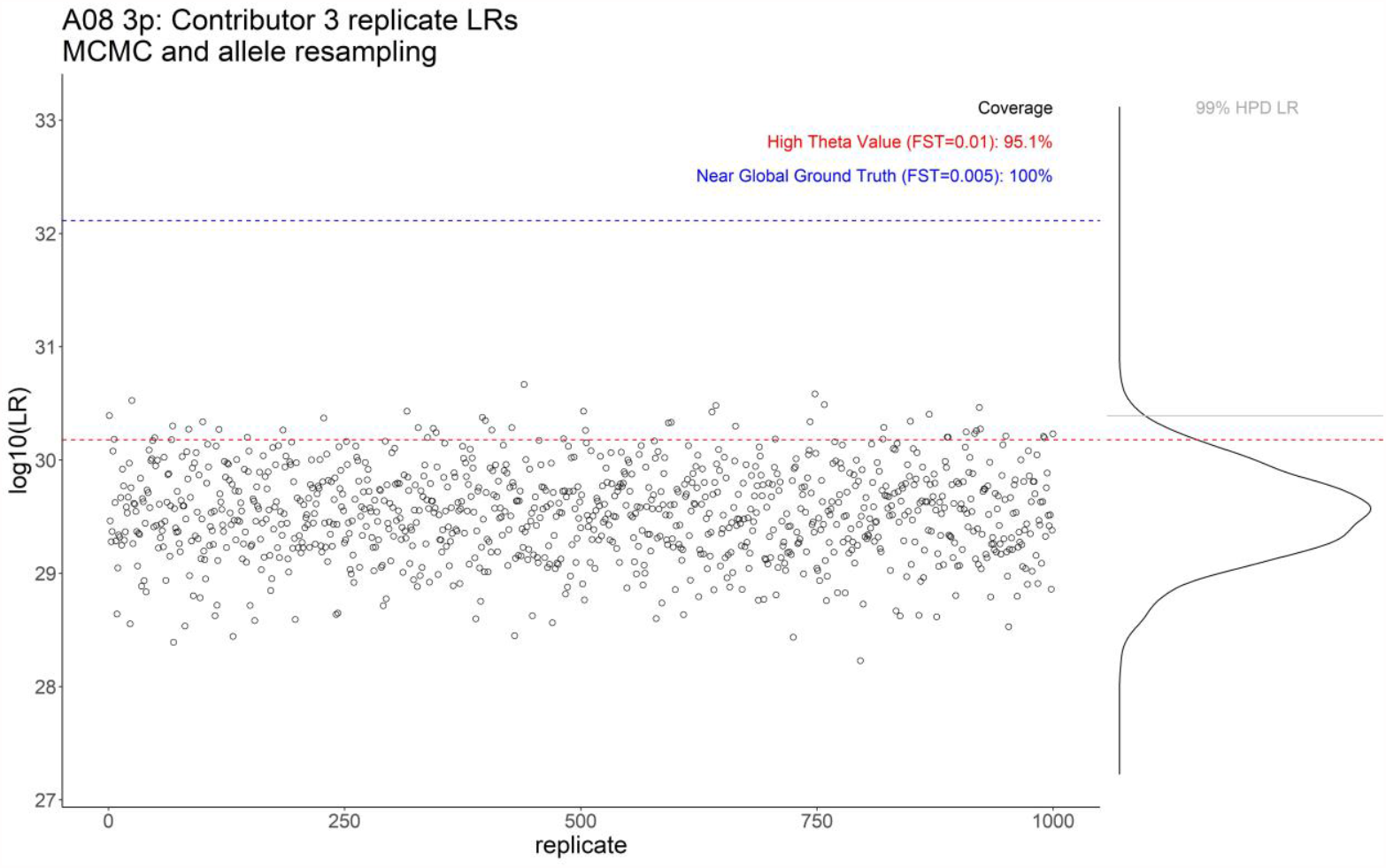
Comparison of each of 1000 replicates for the sample A08 3p contributor reference 3 comparison. The individual log_10_*LR* (α = 99% lower bound inclusive of both MCMC and allele probability resampling, with *θ* = 0.01) are given as hollow black circles. Also shown are HTV as a red dashed line and NGGT as a blue dashed line. At the right-hand margin these data are given as a probability density distribution, and the upper 99% quantile of the HPD density distribution is marked as a grey line on the distribution. The percentage coverage with respect to each ground truth *LR* is provided in the legend.

A review of the range of *LR*s for all comparisons showed high variability in the replicate HPD *LR*s for sample sim 2p Contributor 1 and 1p excl. The variability within the central 95% of sub-source point estimate *LR*s (with *θ* =0.01) was found to be within the expected one order of magnitude for all samples except sim 2p Contributor 1, sim 3p Contributor 1, 2p excl, and 1p excl.

Contributor 1 to sim 2p gave the highest *LR* in the major contributor position for all but four replicates, where the contributor order giving the highest *LR* changed to the minor contributor position. These are the four lowest datapoints around log_10_*LR*∼7. The mixture is a low-level profile simulated with templates of 310 rfu and 61 rfu for each contributor. The known reference profile for contributor 1 had been revised at locus D3S1358 to be a poor fit to the profile, with a corresponding genotype weight of around 10^−5^ in order to force an *LR*<1 at this locus. In the four outlier deconvolutions, this genotype was not accepted for the major contributor.

Samples sim 2p, sim 3p, 2p excl and 1p excl were reinterpreted 1000 times each with extended MCMC accepts (10× the default to give 100,000 burn-in accepts and 500,000 post burn-in accepts for each of the eight chains). Increasing the number of accepts has previously been shown to reduce the variation in the *LR* between replicate interpretations of the same evidence [14]. This is also true for these four samples which showed significant reduction in the variability of the *LR* after running with extended accepts. The central 95% of sub-source *LR*s (with *θ* =0.01) were within one order of magnitude of each other for all four comparisons. All deconvolutions of sim 2p with extended accepts resulted in Contributor 1 aligning with the major contributor position (Figure 5).

**Figure 5.**
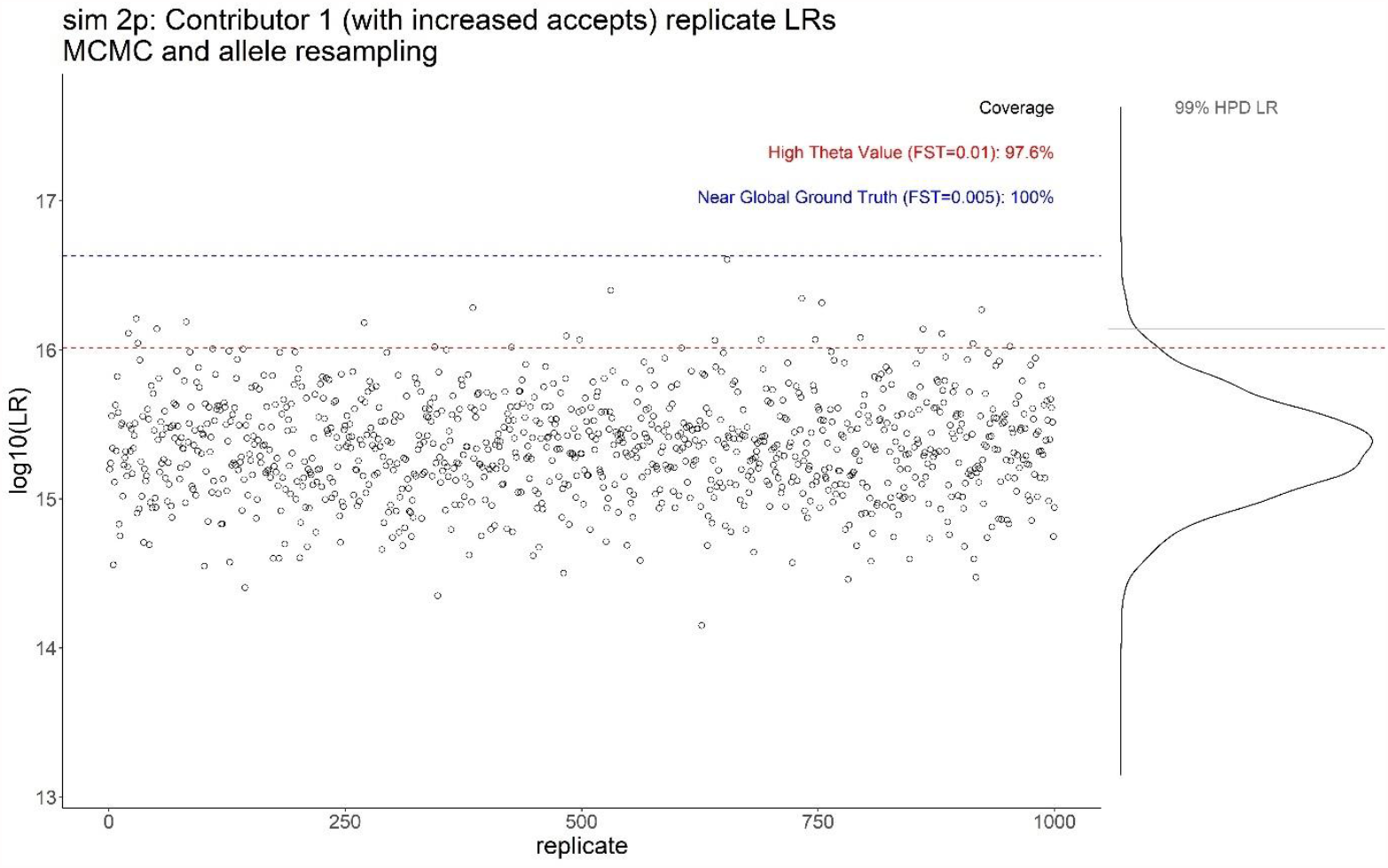
Comparison of each of 1000 replicates with increased MCMC accepts (10× default) for the sample sim 2p Contributor 1.

## Discussion

It is well known that the highest posterior density interval (HPD) of an estimate is not obtained by a function of the HPDs of the parameters. Hence it is not necessary, or even desirable, for each parameter to give 99% coverage as we are interested only in the coverage of the final estimate, in this case the *LR*.

Examination of Table 1 suggests that the near global coverage for all tested samples taking into account MCMC uncertainty, allele probability uncertainty, and subpopulation effects (*θ* = 0.005)was greater than 99% for inclusionary average *LR*s of known donors. The lowest coverage was greater than 95% for an exclusionary *LR* (2p excl). The minimum coverage considering both MCMC and allele probability uncertainty when a typical conservative theta value was applied (*θ* = 0.01) was 92.8%, with the average being around 96%. This suggests that when included in the probability interval method within STRmix^™^ the other layers of conservativeness adequately compensate for the intermittent underperformance of the MCMC variability component.

Based on the data tested, when HPD is applied inclusive of both MCMC and allele frequency uncertainty 99% coverage is reached with respect to our best estimate of the ground truth *LR* (NGC) for inclusionary *LR*s.

Individuals that are poor fits to the profile data were shown to result in more variable *LR*s. The reproducibility of *LR*s between replicate interpretations was improved by increasing the number of MCMC accepts and HPD *LR*s were shown to be generally within one or two orders of magnitude of each other. Increasing the number of accepts also reduced the variability in the point *LR*s for the sim 2p Contributor 1 and 1p excl comparisons. The central 95% of sub-source *LR*s (with *θ* =0.01) were within one order of magnitude of each other for all comparisons with either the default or a 10-fold increase in the number of accepts (data not shown).

We do not want this discussion to give the impression that the lower bound is the expression of the value of the evidence. Pedantically it is a number at the lower end of the plausible range for the value of the evidence. It is as wrong to substitute the lower bound for the mean value as it is to say that all Americans are as short as the shortest American found.

An entire special issue of Science and Justice was dedicated to discussing the topic of precision in *LR*s (we point the reader to Science & Justice 2016, volume 56, issue 5, [15-22], preceded by a discussion in Law, Probability & Risk [23, 24]) and many insightful points were made which we do not recount here.

## Acknowledgements

This work was supported in part by grant 2020-DQ-BX-0022 from the US National Institute of Justice. Points of view in this document are those of the authors and do not necessarily represent the official position or policies of their organisations.

## Appendix Comparison plots

Comparison of each of 1000 replicates for the samples and donors displayed in Table 2. The individual log_10_*LR* (*α* = 99% lower bound inclusive of both MCMC and allele probability resampling, with *θ* = 0.01) are given as hollow black circles. Also shown are HTV as a red dashed line and NGGT as a blue dashed line. At the right-hand margin these data are given as a probability density distribution, and the upper 99% quantile of the HPD density distribution is marked as a grey line on the distribution. The percentage coverage with respect to each ground truth *LR* is provided in the legend.

**Table 2:** A comparison of the coverage test results where only MCMC uncertainty was included [1] 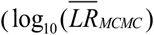 and MCMC coverage) and where the test was inclusive of both MCMC and allele probability resampling, and conservative value for theta, NGGT (*θ* = 0.01) and NGC (*θ* = 0.005).

| Sample | Contributor | $\log_{10}(\overline{LR}_{MCMC})$<br>$\theta = 0.01$ | MCMC Coverage | NGGT | MCMC and allele<br>probability coverage<br>using $\theta = 0.01$ | MCMC and allele<br>probability coverage<br>using $\theta = 0.005$ (NGC) |
| --- | --- | --- | --- | --- | --- | --- |
| A02 2p | 1 | 6.5 | 93.8% | 6.9 | 97.6% | 100% |
|  | 2 | 5.8 | 98.1% | 6.4 | 98.3% | 100% |
| A05 2p | 1<sup>^</sup> | N/A |  |  |  |  |
|  | 2 | 30.9 | 81.7% | 32.2 | 92.8% | 100% |
| A08 3p | 1 | 8.0 | 88.8% | 8.4 | 94.4% | 99.8% |
|  | 2 | 7.5 | 90.7% | 7.9 | 94.9% | 99.9% |
|  | 3 | 30.2 | 76.1% | 32.1 | 95.1% | 100% |
| C06 A 1p | 1 | 22.8 | 100.0% | 24.5 | 97.7% | 100% |
| C06 B 1p | 1 | 13.4 | 100.0% | 13.9 | 96.9% | 100% |
| C10 3p | 1 | 2.9 | 95.7% | 3.3 | 96.1% | 100% |
|  | 2 | 11.0 | 95.1% | 11.6 | 95.1% | 99.9% |
|  | 3 | 12.3 | 92.3% | 13.3 | 95.9% | 100% |
| E05 2p | 1 | 17.9 | 91.7% | 18.4 | 97.1% | 100% |
|  | 2 | 31.8 | 84.7% | 33.1 | 93.1% | 100% |
| sim 3p | 1 | 10.3 | 91.9% | 11.0 | 93.9% | 99.5% |
|  | 2 | 8.5 | 92.5% | 9.0 | 95.5% | 99.9% |
|  | 3 | 1.3 | 96.4% | 1.4 | 97.8% | 100% |
| sim 2p | 1<sup>%</sup> | 16.0 | 100.0% | 16.6 | 99.7% | 100% |
|  | 2* | 1.6 | 98.6% | 2.0 | 99.0% | 100% |
|  | 3 | 4.1 | 97.8% | 4.5 | 98.2% | 100% |
| 1p excl | 1 | -10.0 | 100.0% | -9.7 | 100% | 100% |
| 2p excl | 1 | -5.8 | 93.1% | -5.8 | 94.7% | 95.9% |

<sup>^</sup> Note that the LR was not assigned for reference 1 to sample A05 2p within the original paper
<sup>%</sup> Note that contributor 1 to the simulated two-person mixture was a revision of the true contributor's profile at locus D3S1358. This was done in order to force an LR less than 1 at that locus
<sup>\*</sup> Note that contributor 2 to the simulated two-person mixture was a revision of the true contributor's profile who we term contributor 3. This was done in order to lower the LR closer to 1

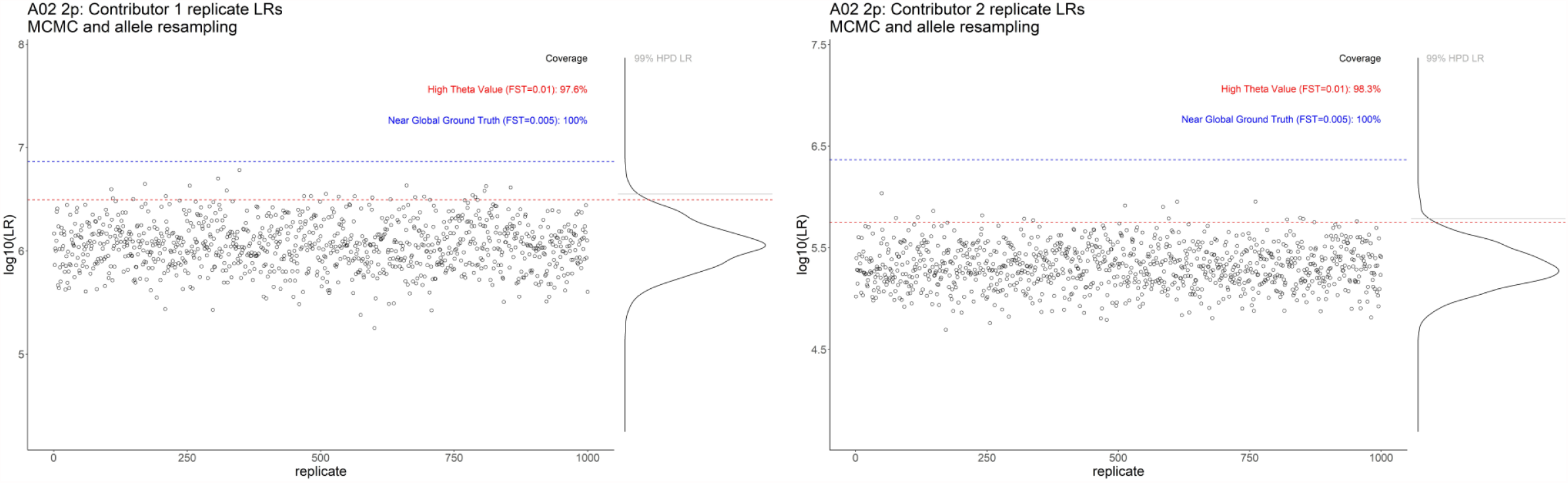
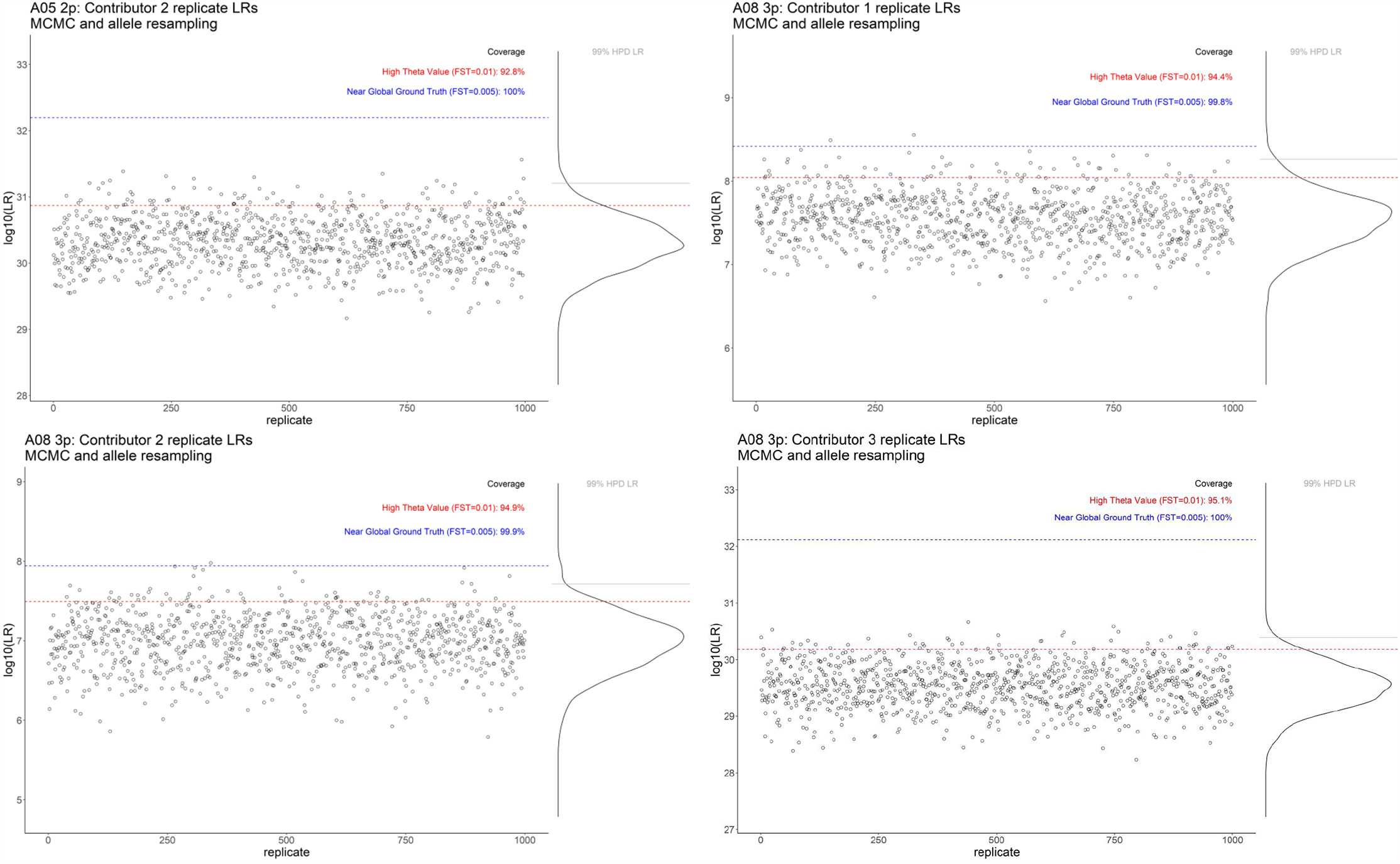
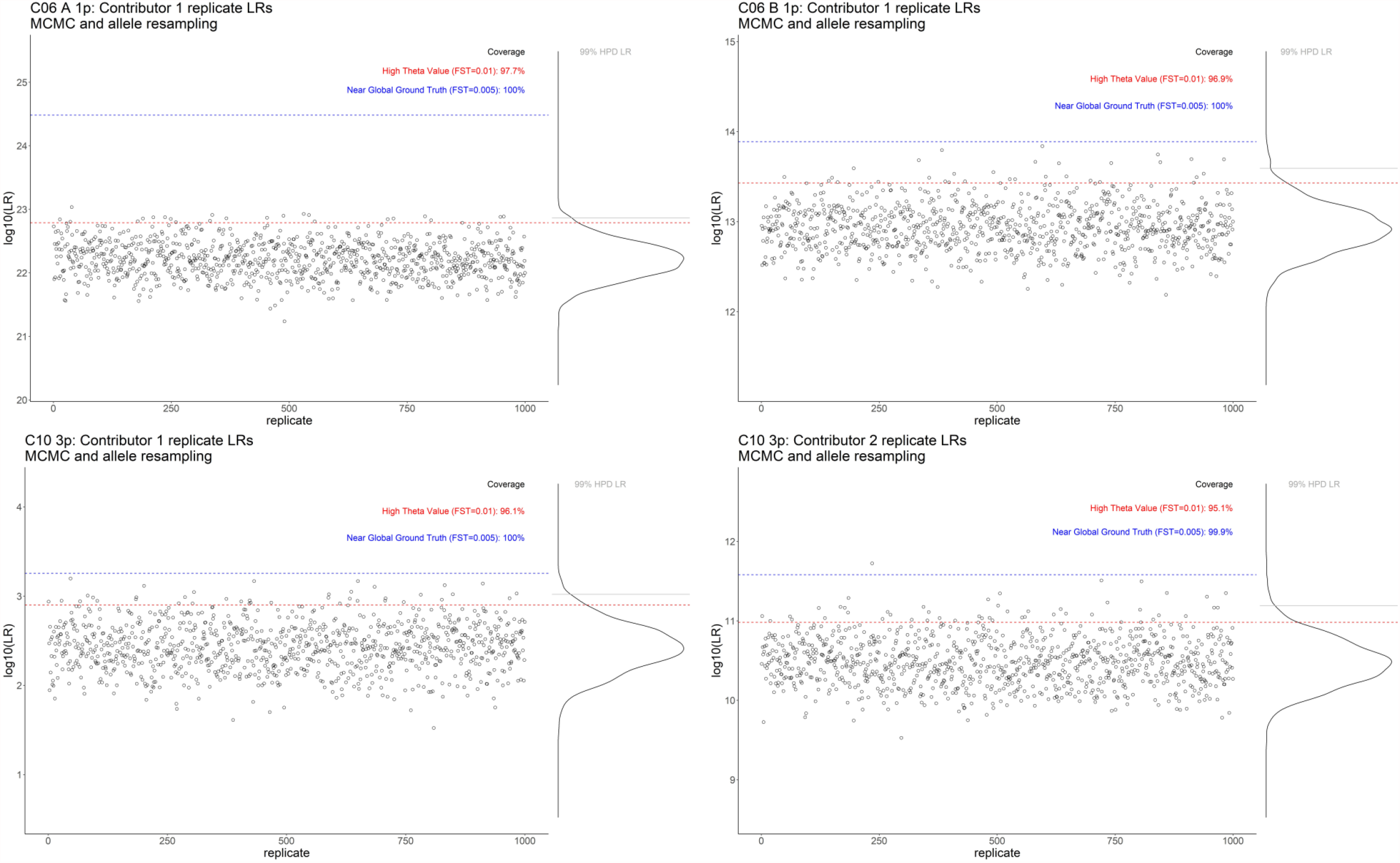
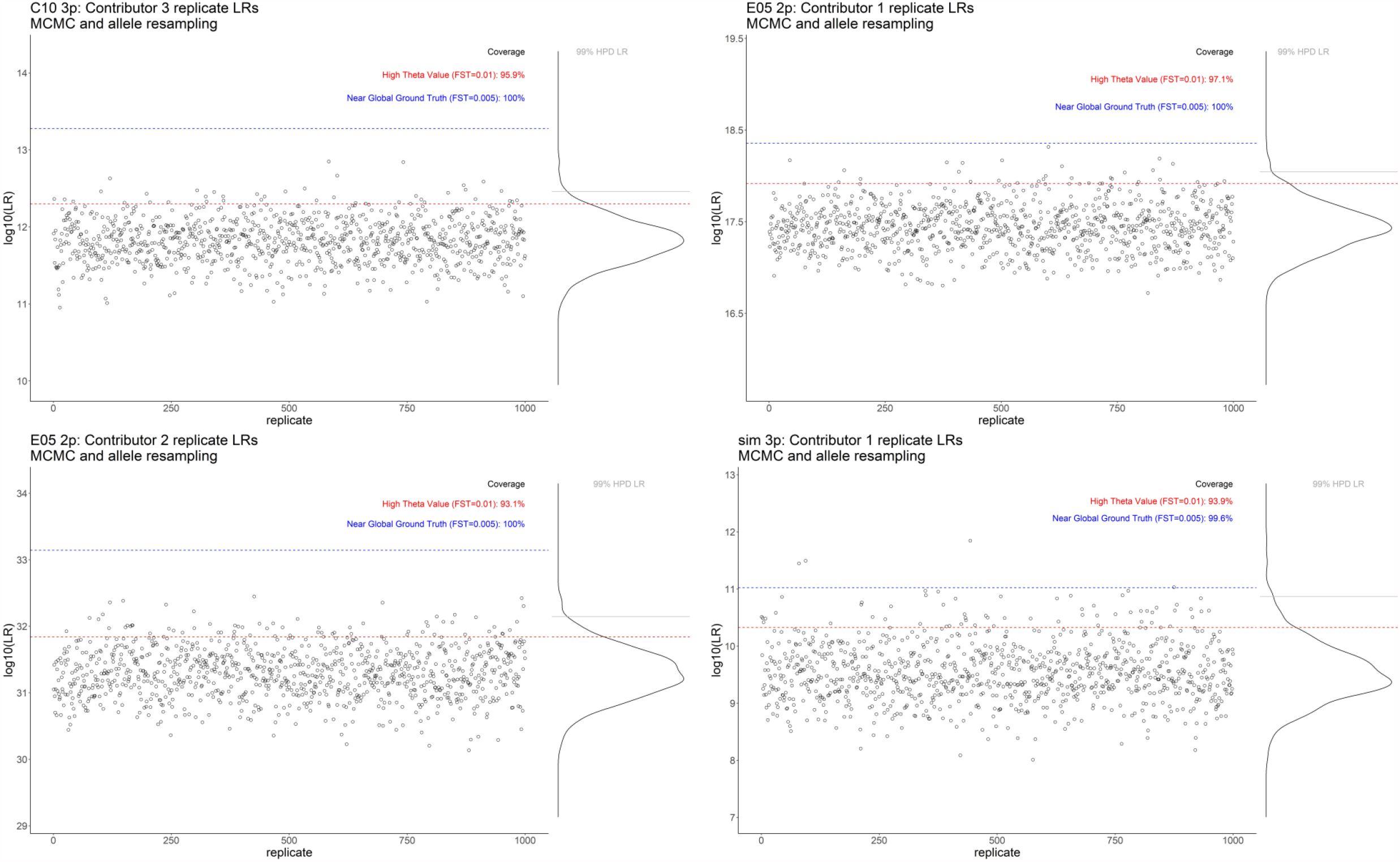
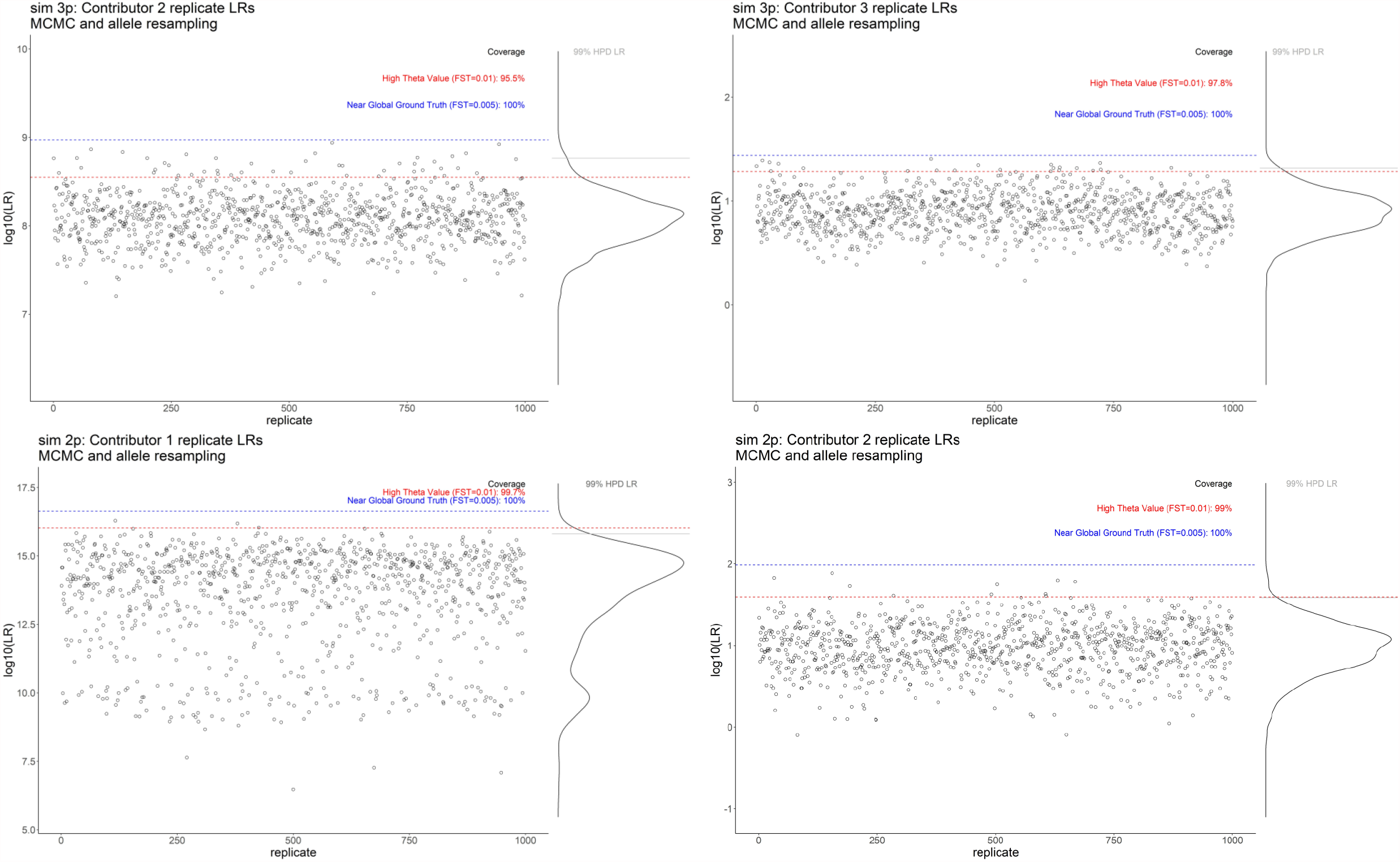
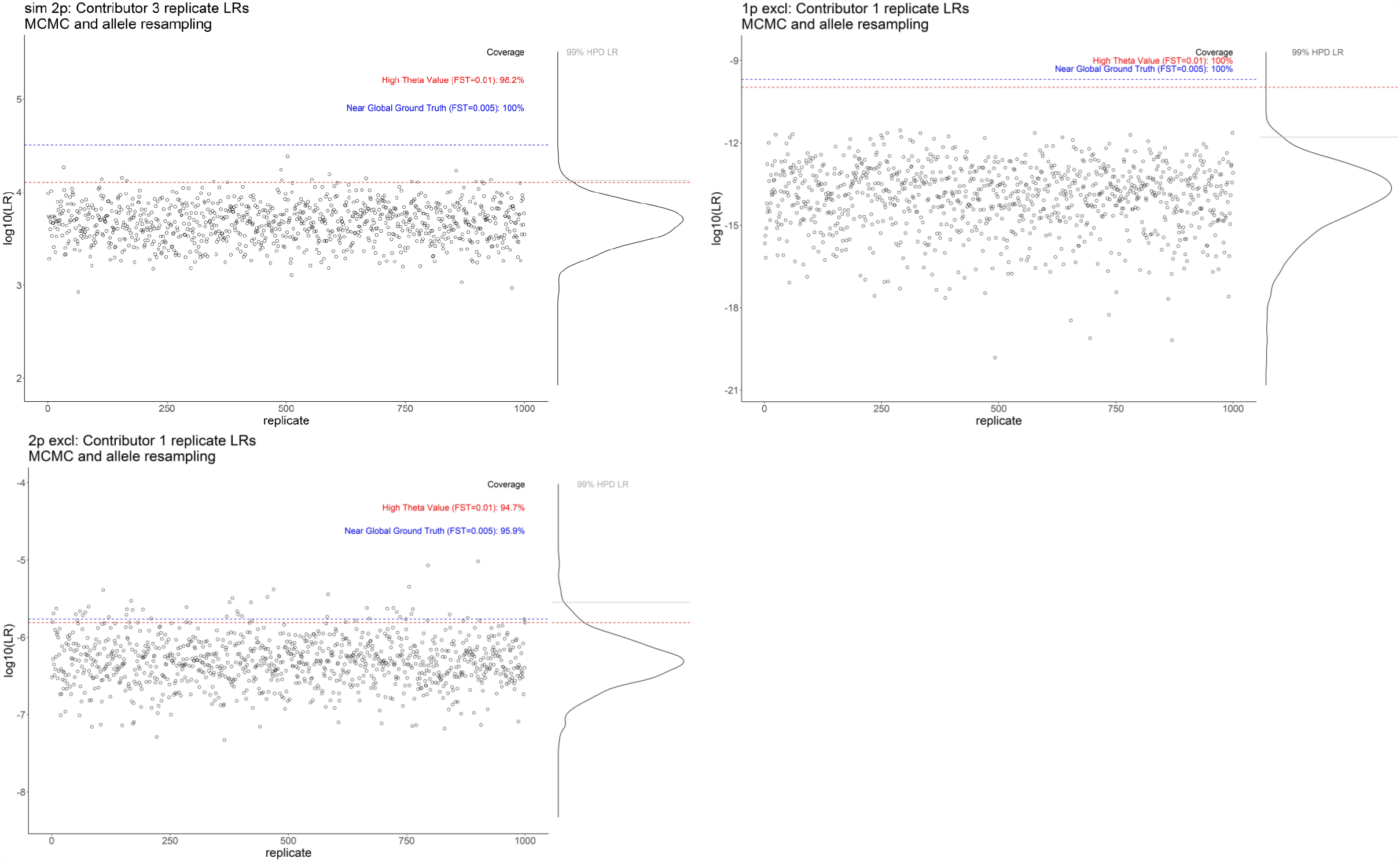

## References

[1] Bright J-A, Taylor D, Curran J, Buckleton JS. Testing methods for quantifying Monte Carlo variation for categorical variables in Probabilistic Genotyping. 2020. https://figshare.com/articles/report/Testing_methods_for_quantifying_Monte_Carlo_variation_for_categorical_variables_in_Probabilistic_Genotyping/13180610. Accessed: 15 March 2021.

[2] Balding DJ. Weight-of-evidence for forensic DNA profiles. Chichester: John Wiley and Sons; 2005.

[3] Kelly H, Bright J-A, Coble MD, Buckleton JS. A description of the likelihood ratios in the probabilistic genotyping software STRmix™. WIREs Forensic Science. 2020;2:e1377.

[4] National Research Council Report: The evaluation of forensic DNA evidence. Washington DC: National Academy Press; 1996.

[5] Balding DJ, Nichols RA. DNA profile match probability calculation: how to allow for population stratification, relatedness, database selection and single bands. Forensic Science International. 1994;64:125–40.

[6] Buckleton J, Curran J, Walsh S. How reliable is the sub-population model in DNA testimony? Forensic Science International. 2006;157:144–8.

[7] Curran JM, Buckleton JS, Triggs CM. What is the magnitude of the subpopulation effect? Forensic Science International. 2003;135:1–8.

[8] Bright J-A, Taylor D, Curran JM, Buckleton JS. Developing allelic and stutter peak height models for a continuous method of DNA interpretation. Forensic Science International: Genetics. 2013;7:296–304.

[9] Taylor D, Bright J-A, Buckleton J. The interpretation of single source and mixed DNA profiles. Forensic Science International: Genetics. 2013;7:516–28.

[10] Alfonse LE, Garrett AD, Lun DS, Duffy KR, Grgicak CM. A large-scale dataset of single and mixed-source short tandem repeat profiles to inform human identification strategies: PROVEDIt. Forensic Science International: Genetics. 2018;32:62–70.

[11] Moretti TR, Moreno LI, Smerick JB, Pignone ML, Hizon R, Buckleton JS, et al. Population data on the expanded CODIS core STR loci for eleven populations of significance for forensic DNA analyses in the United States. Forensic Science International: Genetics. 2016;25:175–81.

[12] Buckleton J, Curran J, Goudet J, Taylor D, Thiery A, Weir BS. Population-specific F_ST_values for forensic STR markers: A worldwide survey. Forensic Science International: Genetics. 2016;23:91–100.

[13] Steele CD, Court DS, Balding DJ. Worldwide FST Estimates Relative to Five Continental Scale Populations. Annals of Human Genetics. 2014;78:468–77.

[14] Bright J-A, Taylor D, McGovern CE, Cooper S, Russell L, Abarno D, et al. Developmental validation of STRmix™, expert software for the interpretation of forensic DNA profiles. Forensic Science International: Genetics. 2016;23:226–39.

[15] Morrison G. Special issue on measuring and reporting the precision of forensic likelihood ratios: Introduction to the debate. Science & Justice. 2016;5:371–3.

[16] Morrison G, Enzinger E. What should a forensic practitioner’s likelihood ratio be? Science & Justice. 2016;5:374–9.

[17] Curran J. Admitting to uncertainty in the LR. Science & Justice. 2016;5:380–2.

[18] Ommen D, Saunders C, Neumann C. An argument against presenting interval quantifications as a surrogate for the value of evidence. Science & Justice. 2016;5:383–7.

[19] Berger C, Slooten K. The LR does not exist. Science & Justice. 2016;5:388–91.

[20] Biedermann A, Bozza S, Taroni F, Aitken C. Reframing the debate: A question of probability, not of likelihood ratio. Science & Justice. 2016;5:392–6.

[21] Hout Avd, Alberink I. Posterior distribution for likelihood ratios in forensic science. Science & Justice. 2016;5:397–401.

[22] Taylor D, Hicks T, Champod C. Using sensitivity analyses in Bayesian networks to highlight the impact of data paucity and direct future analyses: a contribution to the debate on measuring and reporting the precision of likelihood ratios. Science & Justice. 2016;56:402–10.

[23] Nordgaard A. Comment on ‘Dismissal of the illusion of uncertainty on the assessment of a likelihood ratio’ by Taroni F., Bozza S., Biederman A. and Aitken C. Law, Probability and Risk. 2016;15:17–22.

[24] Taroni F, Bozza S, Biedermann A, Aitken C. Dismissal of the illusion of uncertainty in the assessment of a likelihood ratio. Law, Probability and Risk. 2016;15:1–16.

